# Leveraging Sequential and Spatial Neighbors Information by Using CNNs Linked With GCNs for Paratope Prediction

**DOI:** 10.1101/2020.10.15.339168

**Authors:** Shuai Lu, Yuguang Li, Fei Wang, Xiaofei Nan, Shoutao Zhang

**Affiliations:** School of Information Engineering, Zhengzhou University, Zhengzhou, Henan, 450001, China; College of Economics and Management, Zhengzhou University of Light Industry, Zhengzhou, Henan, 450001, China; School of Nursing and Health, Zhengzhou University, Zhengzhou, Henan, 450001, China; School of Life Science, Zhengzhou University, Zhengzhou, Henan, 450001, China

**Keywords:** CNNs, GCNs, attention, paratope prediction

## Abstract

Antibodies consisting of variable and constant regions, are a special type of proteins playing a vital role in immune system of the vertebrate. They have the remarkable ability to bind a large range of diverse antigens with extraordinary affinity and specificity. This malleability of binding makes antibodies an important class of biological drugs and biomarkers. In this article, we propose a method to identify which amino acid residues of an antibody directly interact with its associated antigen based on the features from sequence and structure. Our algorithm uses convolution neural networks (CNNs) linked with graph convolution networks (GCNs) to make use of information from both sequential and spatial neighbors to understand more about the local environment of target amino acid residue. Furthermore, we process the antigen partner of an antibody by employing an attention layer. Our method improves on the state-of-the-art methodology.

## 1 Introduction

ANTIBODY, also known as immunoglobulin, is a Y-shaped protein consisting of two light chains and two heavy chains[1], and can bind to a specific surface of the antigen, named epitope. Amino acid residues of an antibody directly involved in binding epitope is called paratope[2]. The accurate recognition of paratope on a given antibody would greatly improve antibody affinity maturation[3]-[5] and de novo design[6]-[8].

We can get high resolution structure of antibody and antigen complex by experimental methods, such as X-ray[9], NRM[10] and Cryo-EM[11]. However, it remains time consuming and empirical[12]. As more and more protein structures including antibody-antigen complexes have been analyzed, the machine learning-based methods can be used for predicting paratope by learning the paratope-epitope interaction patterns from known antibody-antigen complex structures. According to the type of selecting neighbors of target residue for representing and predicting, the machine learning-based methods can be divided into two categories, leveraging sequential neighbors or spatial neighbors. As for methods leveraging sequential neighbors, a part of the antibody sequence is used consisting of target residue and additional forward and backward sequential neighbors. Sequential neighbors were selected from the whole sequence of antibody like the methods in [13]-[15], and others only took advantage of the sequence of CDR region[16],[17].

Although the sequence is always available at the stages of an antibody discovery campaign earlier than the structure, machine learning-based methods using spatial neighbors can provide more precise definition of the paratope. In [18], the antibody surface patch which was a set of amino acid residues adjacent to each other on the antibody surface, were represented by 3D Zernike descriptors. And the state-of-art method[19] represented an antibody as a graph where each amino acid residue was a node and K nearest spatial neighbors were used in the convolution operator.

In this work, we utilize the sequential and spatial neighbors of target antibody residue by using Convolutional Neural Networks (CNNs) linked with Graph Neural Networks (GCNs) for paratope prediction. Fig.1 shows a diagram of our prediction method and illustrates how the sequential and spatial neighbors information is used to predict binding probability of the target antibody residue. First, we construct an antibody residue feature matrix form sequence-based and structure-based features. Next, we employ CNNs which take the residues feature matrix with a fixed window size as input for considering the influence of sequential neighbors. Then, the output of CNNs are directly fed to GCNs for learning the local environment of spatial neighbors. At last, our program predicts the binding probability of each antibody residue. We also compare results with other existing paratope predictors, and our framework achieves the best performances. Moreover, we add an attention layer to our best performing model attempting to gain more information from antigen partner.

## 2 Materials and Methods

### 2.1 Datasets

We use the datasets the same as [19]. All the complexes in training set are collected by [18] from the training set used to train Paratome[13], Antibody i-Patch[15] and Parapred[16] predictors. The complexes in test set are fetched from AbDb database[20]. The antibody-antigen complexes present in AbDb are split into two categories depending on whether their antigen is a protein or not. In both training and test sets, the complexes whose resolution better than 3.0Å or the antibody sequence which has more than 95% sequence identity are removed. The training set is further split into two disjoint sets: a reduced training set and a validation set, and the validation set is used to tune the hyper parameters in the predictive model.

Structures with nonprotein-binding antibodies are removed in the state-of-art method[19] resulting in 205 complexes for training, 103 for validation and 152 for testing. Specifically, the complexes with PDB ID 2AP2 and 2KVE, only has one chain in antibody which are still retained in this study. Dataset sizes are shown in TABLE 1. Positive residues are residue pairs that participate in the interface, negative residues are pairs that do not. Because in any given complex the size of positive and negative residues is very imbalanced, we use a weighted loss function when training our model.

**TABLE 1.** Number of complexes and residues in the datasets.

| DataSet | Complexes | Positive residues | Negative residues |
| --- | --- | --- | --- |
| Train | 205 | 4449 (5.19%) | 81283 (94.81%) |
| Validation | 103 | 2237 (5.24%) | 60584 (94.76%) |
| Test | 152 | 3314 (5.19%) | 40480 (94.81%) |

### 2.2 Residue Representation

To construct the input matrix, we encode the 1D antibody sequence to a 2D numerical matrix with dimension (*L, N*), where *L* is the length of the antibody sequence and *N* is the residue features vector dimension (128 here).

As shown in Fig.2, the feature representation for amino acid residue *a* is donated by *x*_*a*_. Different components of the feature representation are denoted by the superscripts. Each box indicates the program used to extract a given set of features. All those features can be classified into two classes according to the source: sequence-based and structure-based.

#### 2.2.1 Sequence-based Features

One-hot encoding 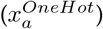: The type of amino acid residue (only 20 possible natural types are considered) is encoded to a 20 dimensional vector, where each element is either 1 or 0 and 1 indicates the existence of a corresponding amino acid residue.

Seven physicochemical parameters 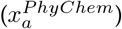 Those parameters are about physicochemical properties of residues summarized by [21].

Profile features 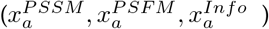: We run PSI-BLAST[22] against the nonredundant (nr)[23] database for every antibody sequence. Then we get the PSSM and PSFM matrix, both with dimension (*L*, 20), as well as a 1D vector related with column entropy with dimension *L*, where *L* is the length of the antibody sequence.

#### 2.2.2 Structure-based Features

Relative accessible surface area 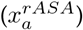, Secondary structure 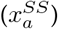, Phi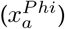 and Psi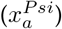 torsion angles for each residue: Those features are computed using DSSP[24]. The secondary structure totally has eight classes and is represented by one-hot encoding.

Half sphere amino acid composition 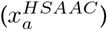: HSAAC captures the amino acid residue composition in the direction of the side chain of a residue, defined as the number of times a particular amino acid occurs in that direction within a minimum atomic distance threshold of 8.0Å from the residue of interest.

Residue depth 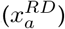: We calculate the average distance of the atoms of a residue from the solvent accessible surface by MSMS[25].

Protrusion Index 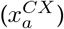: The protrusion index of a non-hydrogen atom is calculated using PSAIA[26] which is defined as the proportion of the volume of a sphere with a radius of 10.0Å centered at that atom that is not filled with atoms[27]. Each element of this vector is normalized to have the range from 0 to 1 as in [28].

B-factor 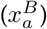: The B-factor (or temperature factor) is an indicator of thermal motion about an atom. We use the maximum B-factor of any atom for each residue.

### 2.3 Antibody Representation and Paratope Definition

We represent an antibody as a graph[19], where each residue is a node whose features represent the properties of the residue. We define the spatial neighbors of a residue as a set of *K* (20, in our work) closest residues determined by the mean distance between their heavy atoms [24]. Fig.3 shows sequential and spatial neighbors of a target residue.

From the analyzed 3D structure of an antibody and antigen complex, a residue on antibody is judged to belong to the paratope if at least one of its heavy atoms is located within 4.5Å from any antigen atoms like previous methods[11],[12].

### 2.4 Convolutional Neural Networks (CNNs) for Processing Sequential Neighbors

The sequence of the input antibody with length *L* is considered as a set of sequential nodes *S* and each node is represented as a 1D vector *s*_*i*_, where 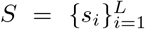. All the nodes of the antibody sequence compose a 2D features matrix as said in Section 2.2.

In order to leverage sequential neighbors information of target residue, we consider a window of fixed size in sequence centered around target residue and concatenate their features as input which can be shown as *s*_*i−w*:*i*+*w*_, where *i* is the index of target residue and the fixed window size=11 (*w* = 5). Before the first and after the last residue of the antibody sequence, we use a default zero padding. Our CNNs all have a fixed stride=1 so that the output *q*_*i*_ will not have dimensional change shown as

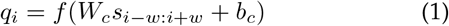

where f is a non-linear activation function (e.g. ReLU), *W*_*c*_ is the wight matrix, and the *b*_*c*_ is the bias vector. Here we use residual connections which act as a shortcut connection between inputs and outputs of some part of a network by adding inputs to outputs which can be shown as

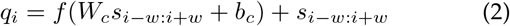

As a result, we apply the function to obtain a set of hidden vector of every position of the antibody sequence: *Q* = {*q*_*i*_ *q*_1_, *q*_2_, *q*_3_, …, *q*_*l*_}

### 2.5 Graph Convolutional Networks (GCNs) for Processing Spatial Neighborhoods

We use the graph convolution[29] which enables aggregation over spatial neighbors of target residue and together contributes to the formation of a binding interface.

For a node *q*_*i*_, the structural environment consisting of *K* spatial neighbors 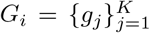 from the input graph, the graph convolution operation results in a vector *z*_*i*_, which can be shown as

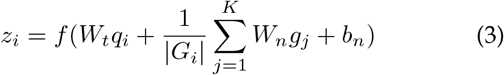

The parameters of this operation include the aggregation weight matrix *W*_*t*_ for target node, the aggregation weight matrix *W*_*n*_ for the neighboring nodes, and the bias vector *b*_*n*_. The dimensionality of the weight matrices is determined by the dimensionality of the inputs and the number of filters.

### 2.6 Classifier

Finally, two fully connected layers perform classification for each antibody residue *z*_*i*_ after processing by CNNs and GCNs. An inverse logit function transforms each residue’s output *y*_*i*_ to indicate the probability of belonging paratope shown as

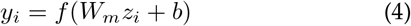

### 2.7 Training Details

We implement our model using PyTorch[30] v1.4. Validation sets are used to find the optimal set of network training parameters for final evaluation. The training details of these neural networks are as follows: optimization: Momentum optimizer with Nesterov accelerated gradients; learning rate: 0.001; batch size: 32; dropout: 0.5; sequential neighbors size: 11 (fixed, including target residue); spatial neighbors in the graph: 20 (fixed); number of layers in GCNs: 1, 2 or 3; number of layers in CNNs: 1, 2 or 3. Training times of each epoch vary from roughly 1-10 minutes depending on network depth, using a single NVIDIA RTX2080 GPU.

For each combination, networks are trained until the performance on the validation set stops improving or for a maximum of 250 epochs. GCNs have the following number of filters for 1, 2 and 3 layers, respectively: (256), (256, 512), (256, 256, 512). All weight matrices are initialized as in [29] and biases are set to zero. Training is carried out by minimizing the weighted cross-entropy loss function[29].

## 3 Results and Discussion

### 3.1 Performances Comparison Between Different Depth Combination of CNNs and GCNs

In this section, we compute precision and recall by predicting residues as paratope with probability above 0.5[19]. As the area under the receiver operating characteristics curve (AUC ROC) is threshold-independent and increases in direct proportion to the overall prediction performance, we take it to assess the overall predictive abilities. Beside, we consider the area under the precision recall curve(AUC PR). To provide robust evaluation of performance, we have trained and tested all networks five times, and computed the mean and standard error.

Results comparing the AUC ROC and AUC PR of various layers combination of CNNs and GCNs are shown in TABLE 2 and TABLE 3. Our first observation is that the all the CNNs linked with GCNs methods, with AUC ROC around 0.97 and AUC-PR around 0.70, outperform the individual CNNs or GCNs methods which have distinct lower AUC PRs, showing that the incorporation of combined information from a residue’s sequential and spatial neighbors improves the accuracy of interface prediction. This matches the biological intuition that the region around a residue should impact its binding affinity[31].

**TABLE 2.** AUC ROC of various layers combination of our networks

| Methods |  | GCN layers |  |  |  |
| --- | --- | --- | --- | --- | --- |
|  |  | 0 | 1 | 2 | 3 |
| CNN layers | 0 | 0.935±0.001 | 0.969±0.001 | 0.973±0.000 | 0.973±0.000 |
|  | 1 | 0.947±0.002 | 0.972±0.001 | **0.975±0.001** | **0.975±0.001** |
|  | 2 | 0.951±0.001 | 0.971±0.000 | 0.973±0.001 | 0.973±0.001 |
|  | 3 | 0.958±0.001 | 0.974±0.001 | **0.975±0.001** | 0.971±0.000 |

**TABLE 3.** AUC PR of various layers combination of our networks

| Methods |  | GCN layers |  |  |  |
| --- | --- | --- | --- | --- | --- |
|  |  | 0 | 1 | 2 | 3 |
| CNN layers | 0 | 0.593±0.006 | 0.687±0.011 | 0.688±0.005 | 0.666±0.002 |
|  | 1 | 0.649±0.010 | 0.703±0.008 | 0.696±0.007 | 0.676±0.005 |
|  | 2 | 0.662±0.004 | 0.700±0.003 | 0.702±0.008 | 0.657±0.008 |
|  | 3 | 0.682±0.003 | 0.705±0.009 | **0.706±0.005** | 0.657±0.005 |

We also observe that the effect of the combination number of CNNs and GCNs layers is not linear, i.e. more layers will not achieve better performance. Indeed, in protein interface prediction, networks with more than four layers performed worse in [29]. In addition, one layer GCN achieves better performance than two layer GCNs about paratope prediction in task-specific learning in [19]. We agree with these findings and draw the same conclusions.

### 3.2 Comparison Between Different Residue Features Combination

As said in Secttion2.2, residue features are classified into two classes: sequence-based and structure-based according to the source. Furthermore, sequence-based features can be divided into three parts: residue type one-hot encoding(a), profile features(b) and the seven physicochemical parameters(c) as their different properties. All the structurebased features are considered as an individual part(d). In order to find the importance of all these residue features, we test different residue features combination on our best model. Because the residue type is the most basic feature, all combination must include it’s one-hot encoding, e.g. a+b, a+c, a+d, a+b+c, a+b+d, a+c+d and a+b+c+d(all).

We obtain the best performance form the model with 3 layers CNNs linked with 2 layers GCNs as shown in TABLE 1 and TABLE 2. Hence, we train this model again using the other 6 kinds of residue features combination. Each combination was evaluated by averaging all the AUC ROC and AUC PR of all antibodies in testing set. Both mean value and standard deviation are reported in Fig.4 and Fig.5.

From Fig. 4, we can see that there are three residue features combination (a+b: 0.968±0.025, a+b+c: 0.953±0.031, a+b+d: 0.969±0.022) almost achieving the optimal performance (0.975±0.019). All of them contain the profile features(b). As for the AUC PR in Fig.5, we can see that performance vary from all kinds of residue features combination. The model using all the features still works best.

**Fig 1.**
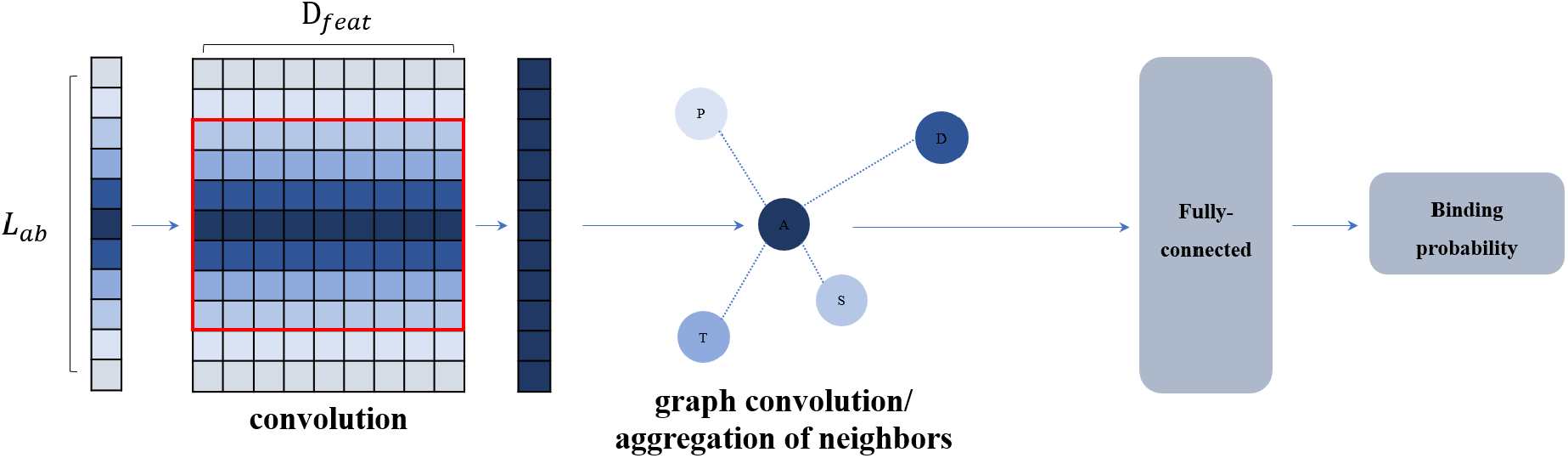
Network architecture. Here, *L*_*ab*_ is the length of antibody sequence and target residue is in deepest blue. The nearer neighbor is in deeper blue. *D*_*feat*_ is the dimension of residue feature vector and a fixed window size of sequential neighbors are within the red square. Our CNNs take a (*L*_*ab*_, *D*_*feat*_) matrix as input and our GCNs use their final output as input. The GCNs make an aggregation of the spatial neighbors. Then the fully connected networks are fed by the output of the final GCN and predict the binding probability for each antibody residue.

**Fig 2.**
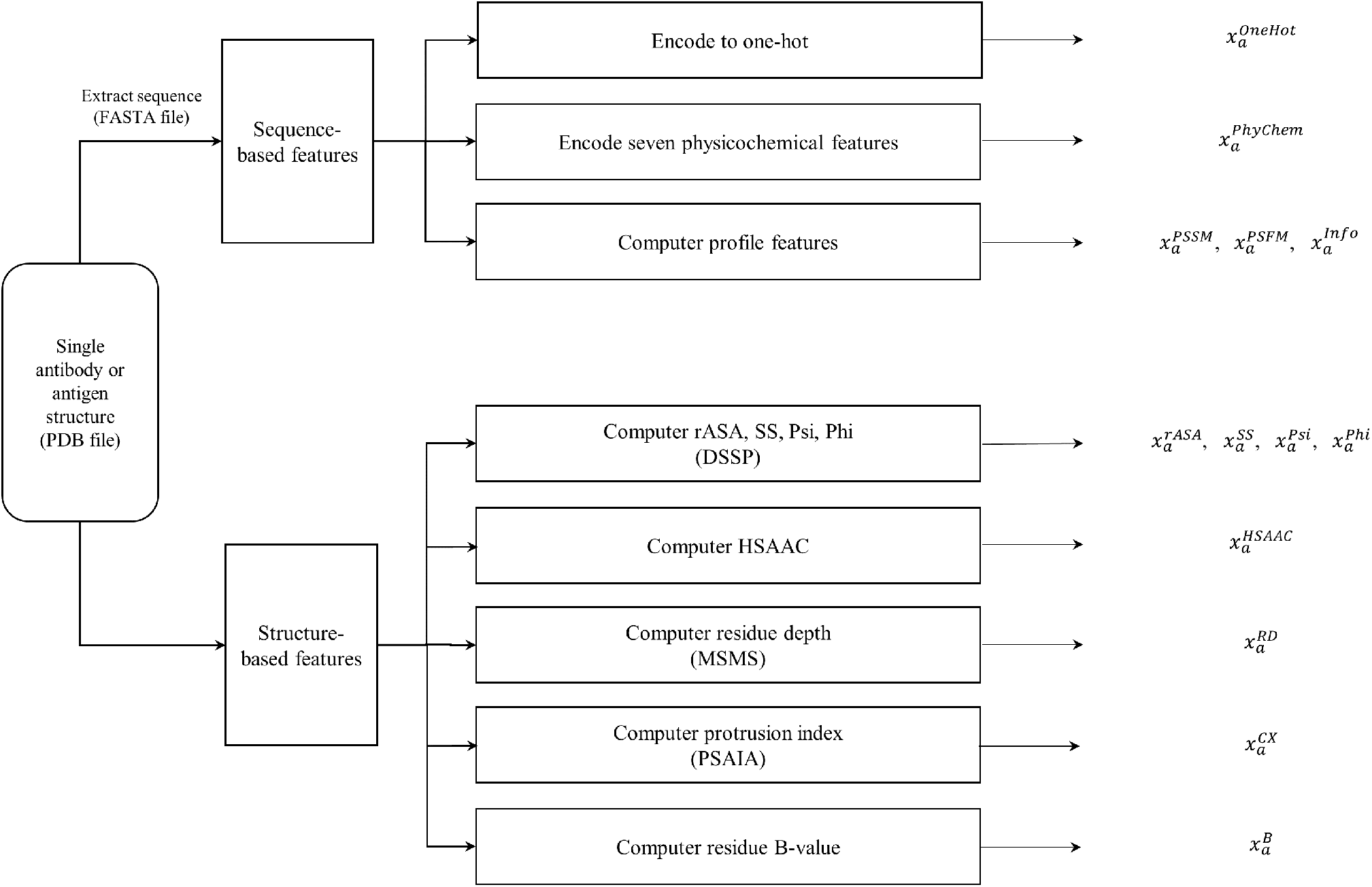
Residue-level feature extraction in this study.

**Fig 3.**
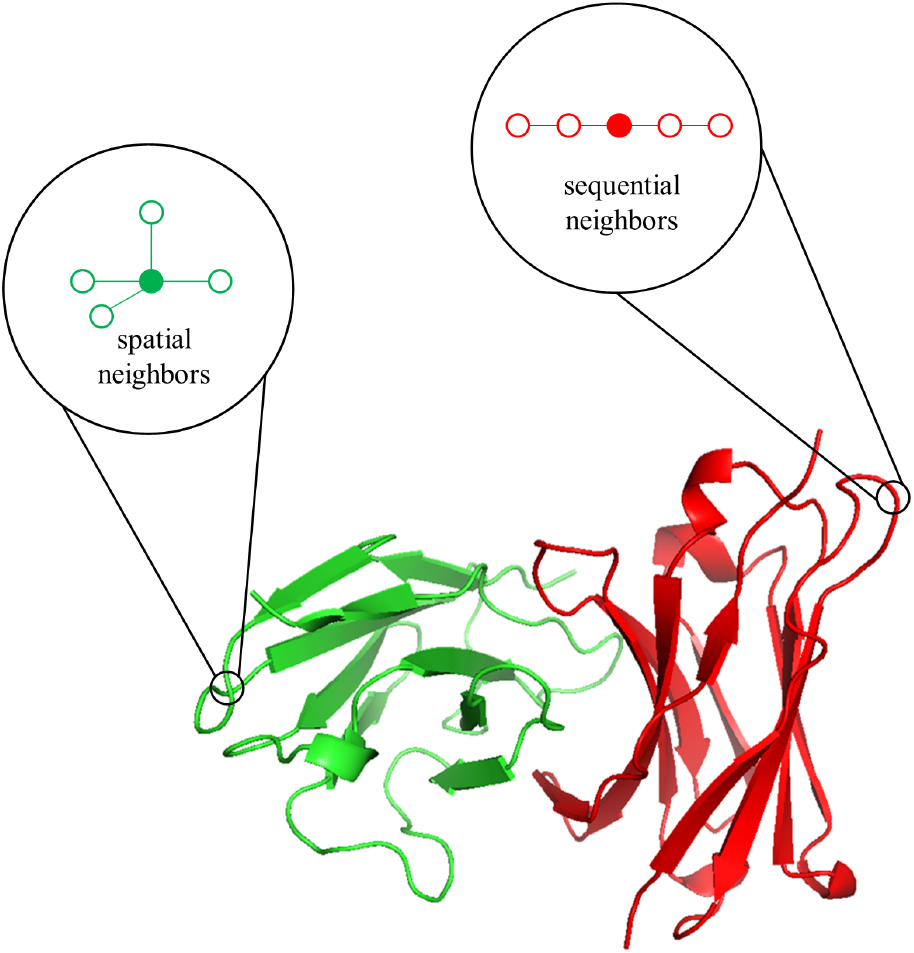
Sequential and spatial neighbors of a target residue(PDB ID: 1A2Y)

**Fig 4.**
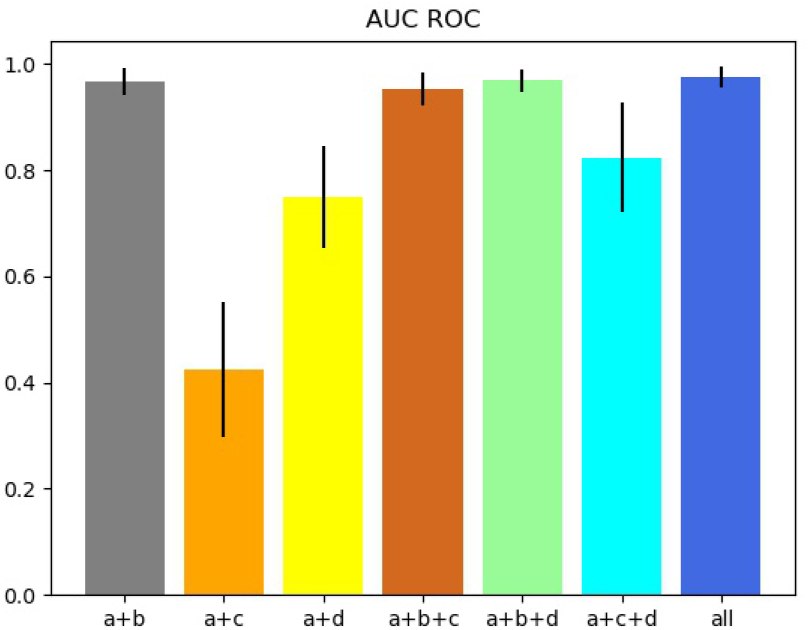
AUC ROC between different residue features combination.

**Fig 5.**
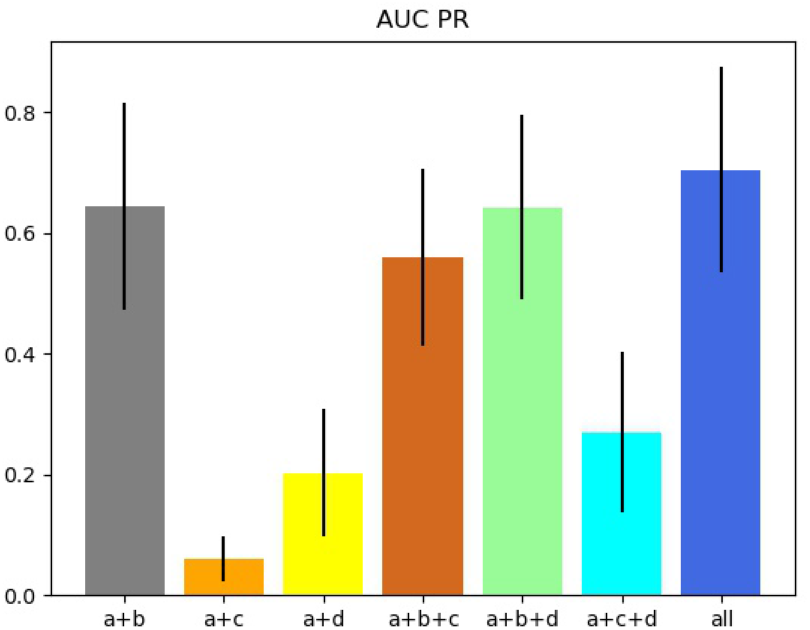
AUC PR between different residue features combination.

### 3.3 Comparison With Existing Predictors of Paratope Prediction

As shown in Fig. 6. and Fig. 7., we compare our method to other existing methods specifically for paratope prediction, i.e. Antibody i-path which pays attention to energetic importance(AUC ROC:0.840, AUC PR: 0.376)[15], Parapred which consists of CNN and RNN-based networks(AUC ROC:0.933, AUC PR: 0.622)[16], model using 3D Zernike descriptors(AUC ROC:0.950, AUC PR: 0.658)[18] and model taking advantage of graph convolution and attention mechanism(AUC ROC:0.958, AUC PR: 0.703)[19].

**Fig 6.**
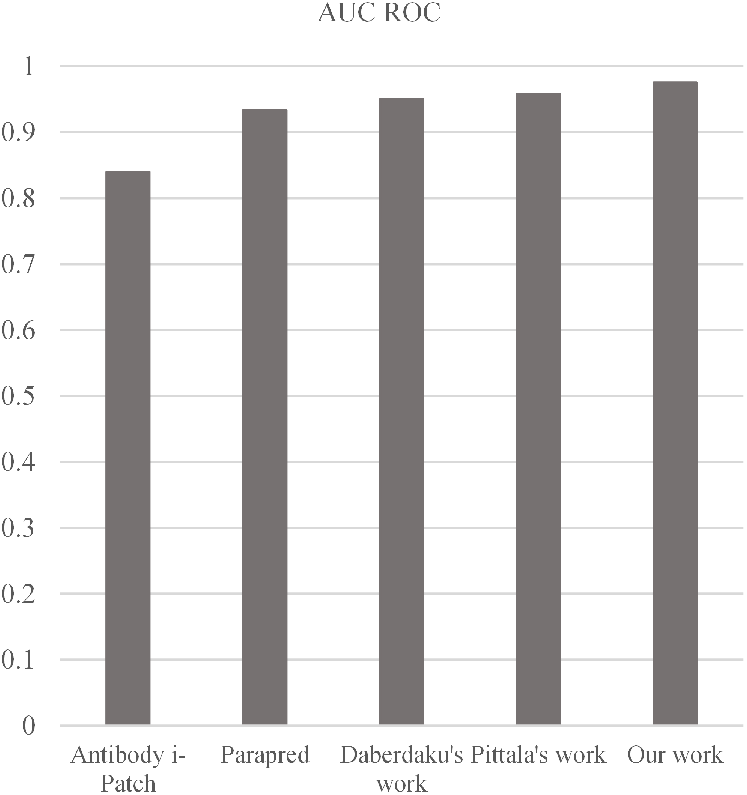
AUC ROC between existing predictors of paratope prediction.

**Fig 7.**
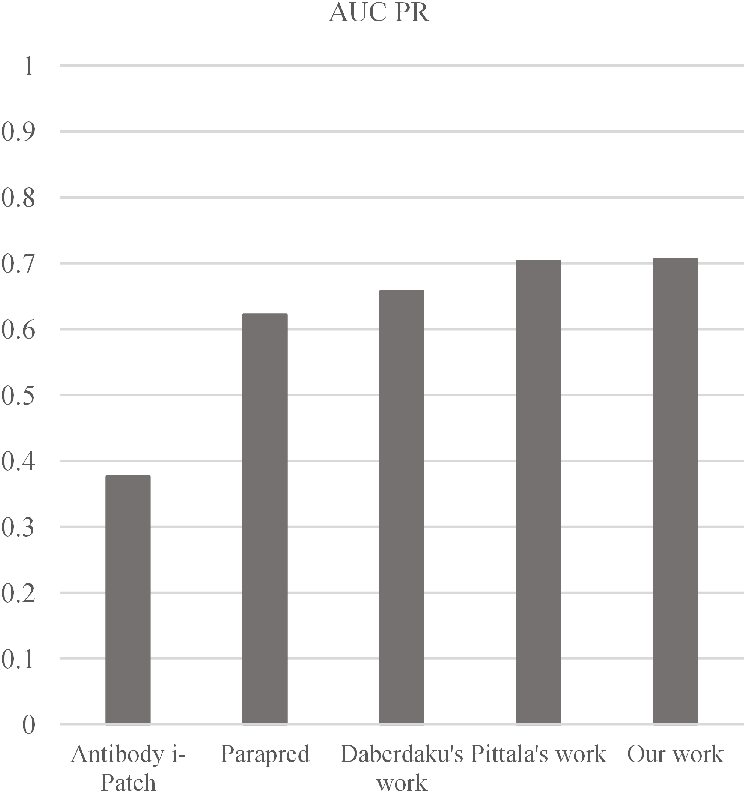
AUC PR between existing predictors of paratope prediction.

**Fig 8.**
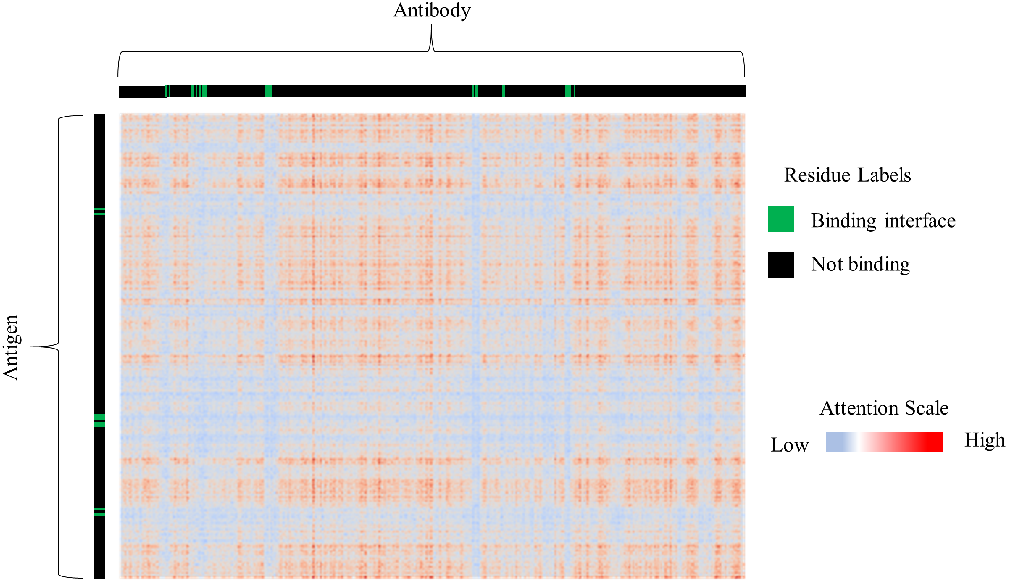
Attention visualization

Note that these methods only considering sequential or spatial neighbors of target antibody residue. Our model achieves greater performance compared to these methods on both AUC ROC(0.975±0.001) and AUC PR(0.706±0.005).

### 3.4 Adding Attention Layer for Processing Antigen Partner

An attention layer was used to explore the specific interaction between antibody and antigen pairs on paratope and epitope prediction[19]. In [19], epitopes had a distinct attention profile compared to other residues on the antigen and the paratope prediction networks perform significantly better at predicting epitopes in the cross-task evaluation. In this study, we add an attention layer the same as [19] to our best model after the GCNs which take both antibody and antigen sequence as input and share the same parameter but resulting in lower performance (AUC ROC: 0.974±0.001, AUC PR:0.698±0.007) for paratope prediction.

Fig.8 shows the heatmap of attention score between every pairs of residues from the complex on which our model perform best (PDB ID 5K59). But we canot see outstanding performance as in epitope predictor, which could be caused by the different environment components of epitope and paratope[15], [34].

## 4 Conclusion

In this study, we design and implement a new structure-based paratope predictor leveraging sequential and spatial neighbors of target antibody residue. Our model is trained on the antibody-antigen complex structures collected from datasets of some paratope predictors which includes the most structures. Moreover, we utilize more residue features consisting of sequence-based and structure-based. Experimental results with a training dataset and an independent validation dataset demonstrate the efficiency of our method.

The superior performances of our method are due to several reasons, including a rich dataset, more sufficient features selection, and careful construction of the prediction model considering sequential and spatial neighbors at same time.

We note that our program has two potential disadvantages. First, the predictor needs antibody structure as it takes structure-based residue features as input. Second, at the stage of extracting residue features, it consumes long computer time as PSI-BLAST[22] needs to be performed. In our future work, we will take adjacent information from antibody sequence so that the predictor can make use of GCNs without structure. We will attempt to accelerate the computation speed by using several servers to concurrently perform PSI-BLAST[22].

Besides, an attention layer improves the performance on epitope prediction but results in a lower in our study due to different environment components of epitope and paratope[15], [34]. In the future, we will try to design a better attention score function for paratope prediction.

Biomolecule binding motifs mining is a long-term challenge for understanding their function. The forming of incorrect interaction between some critical molecules has been revealed as one of the important causes for diseases like COVID-19[32]. The method proposed in this study is specifically for identifying the antibody-antigen binding residues. In the future work, we will further investigate the applicability of our model to other types of molecules binding residues prediction problem, e.g., drug-target interaction prediction[33].

## Acknowledgements

Acknowledgments

This work was supported by the ‘Created Major New Drugs’ of Major National Science and Technology (No. 2019ZX09301-159), and Leading Talents Fund in Science and Technology Innovation in Henan Province(194200510002). Xiaofei Nan and Shoutao Zhang are the corresponding authors for this paper.

